# Closed-loop recruitment of striatal parvalbumin interneurons prevents the onset of compulsive behaviours

**DOI:** 10.1101/2022.01.10.475745

**Authors:** Sirenia Lizbeth Mondragón-González, Christiane Schreiweis C, Eric Burguière E

## Abstract

A prominent electrophysiological feature of compulsive behaviours is striatal hyperactivity, yet, its underlying regulatory processes still need to be characterised. Within the striatum, parvalbumin-positive interneurons (PVI) exert a powerful feed-forward inhibition essential for the regulation of striatal activity and are implied in the suppression of prepotent inappropriate actions. To investigate the potential role of striatal PVI in regulating striatal activity and compulsive behaviours, we used the Sapap3 knockout mice (Sapap3-KO), which exhibit compulsive-like self-grooming. We first showed that the number of compulsive-like events in Sapap3-KO mice was reduced to normal levels by continuous optogenetic activation of striatal PVI in the centromedial striatum. To narrow down the critical time window of striatal PVI recruitment for regulating compulsive-like grooming, we then developed a novel closed-loop optogenetic stimulation pipeline. Upon a predictive biomarker of grooming onsets, characterised by a transient power increase of 1-4 Hz frequency band in the orbitofrontal cortex, we provided real-time closed-loop stimulation of striatal PVI. This targeted closed-loop optogenetics approach reduced grooming events as efficiently as continuous recruitment of striatal PVI with a reduction of stimulation time of 87%. Together, these results demonstrated that recruitment of striatal PVI at the initiation of the compulsive events is sufficient to drastically reduce compulsive-like behaviours and pave the way for targeted closed-loop therapeutic protocols.

## INTRODUCTION

Compulsive behaviours consist of pathological repetitive behaviours executed despite negative consequences. They are a core feature of various neuropsychiatric disorders, including obsessive-compulsive disorder (OCD). Increasing evidence in studies of neuropsychiatric disorders with repetitive behaviours points toward malfunctioning cortico-basal ganglia circuits, which are key players in the formation and regulation of actions^1,2^. In particular, altered fronto-striatal circuits – including the orbito-frontal cortex (OFC) and its primary input site, the centromedial striatum (CMS) – have been observed in OCD patients^3,4^, as well as in rodent models expressing pathological repetitive behaviours^5–7^. Striatal hyperactivity has surfaced as one of the prominent physiological characteristics during compulsive episodes^6–9^, and the reduction of striatal activity correlates with symptom alleviation in OCD patients^10,11^. However, the neuronal abnormalities that underlie this striatal hyperactivity in compulsive behaviours remain to be characterised. Under normal conditions, fast-spiking, parvalbumin-expressing striatal interneurons (PVI) receive strong afferents from the cortex and regulate the activity of striatal medium spiny neurons (MSN) through a powerful feed-forward inhibition, given their earlier and low activation threshold relative to MSN^10,12–15^. These physiological properties have been proposed to adapt and regulate behavioural output via the orchestration and tuning of MSN activity^16–18^. Interestingly, post-mortem studies in patients and mouse models suffering from pathological repetitive behaviours have reported a consistent decrease in PVI density in medial striatal areas^19–22^. This previous evidence suggests a crucial link between decreased PVI function and pathological repetitive behaviours, but a causal link has been lacking to date. To study the contributions of PVI in regulating compulsive behaviours, we used the Sapap3-KO mouse model that displays both striatal hyperactivity and a decreased density of PVI in medial striatal areas^6,23^. To prove and temporally precisely define their role in the emergence of transient and spontaneous compulsive-like behaviours, we have established a closed-loop optogenetic approach with a real-time responsive system, which was triggered by the detection of a reliable, symptom-predictive biomarker.

## RESULTS

### I. Recruitment of striatal parvalbumin-expressing interneurons alleviates compulsive-like behaviours

We bilaterally injected Sapap3^-/-^ :: PV^Cre/Wt^ (Sapap3-KO/PVCre) mice (n=10) with a Cre-dependent adeno-associated viral vector expressing channelrhodopsin-2 (AAV5-hChR2(H134R)-mCherry) into the centromedial striatum (CMS) (Fig. 1a,b). Bilateral implantation of optic fibers at the injection site allowed us to selectively recruit parvalbumin-positive fast-spiking interneurons (PVI) in the CMS (Fig. 1c, Extended Data Fig.1a) via a customised implant designed for simultaneous optogenetic neuromodulation and *in-vivo* electrophysiology (Extended Data Fig. 2). We videotaped naïve freely-moving Sapap3-KO/PVCre mice on three separate days while delivering optogenetic stimulation (5-ms pulses at 20 Hz, 10mW). The experimental paradigm consisted of a 10–15-minute habituation period, followed by ten alternating trials of 3 minutes of active blue light stimulation (ON) and 3 minutes without blue light stimulation (OFF, Fig. 1d). During the PVI optogenetic activation periods, the Sapap3-KO/PVCre mice exhibited a significant decrease of grooming bouts by 55.8% and of grooming duration by 46.25% (Wilcoxon matched-pairs signed-rank tests, *P*_*On-OFF, onsets*_ = 0.004, *P*_*On-OFF, duration*_ = 0.002; GLMM, *P*_*On-OFF, onsets*_ < 0.01, *P*_*On-OFF, duration*_ < 0.0001), to levels comparable to wildtype baselines (Mann Whitney test, *P*_*onsets*_ = 0.93, *P*_*duration*_ = 0.78) (Fig. 1e, f). This effect was consistent across trials (Fig. 1e, f upper panels) and sessions (Fig. 1e, f lower panels, Extended Data Fig. 3). On the contrary, under the same protocol, age-matched control groups including Sapap3-KO/PVCre mice expressing a virus with a fluorescent reporter only (AAV5-mCherry) (n=6) and wildtypes (Sapap3^+/+^:: PV^Cre/wt^) expressing hChR2-mCherry (n=5) did not significantly decrease their grooming activity during the blue light stimulation trials (Wilcoxon matched-pairs signed-rank test: Sapap3-KO/PVCre expressing mCherry: *P*_*On-OFF, onsets*_ = 1, *P*_*On-OFF, duration*_ = 0.31; wildtype mice expressing hChR2-mCherry: *P*_*On-OFF, onsets*_ = 0.81, *P*_*On-OFF, duration*_ = 0.81) (Fig. 1e, f). We looked at individual grooming bouts to further understand the reduction of grooming activity upon optogenetic excitation of striatal PVI in Sapap3-KO/PVCre expressing hChR2-mCherry. We found that the average duration of individual grooming events was consistent between ON and OFF trials (Wilcoxon matched-pairs signed-rank tests, *P* = 0.21) (Fig. 1g), pointing out that the overall decrease in grooming duration was due to a reduced number of grooming bout initiations. Our findings showed that selective stimulation of striatal PVI activity alleviates excessive self-grooming by reducing the number of grooming onsets, suggesting their implication in the regulation of grooming initiations.

**Figure1.**
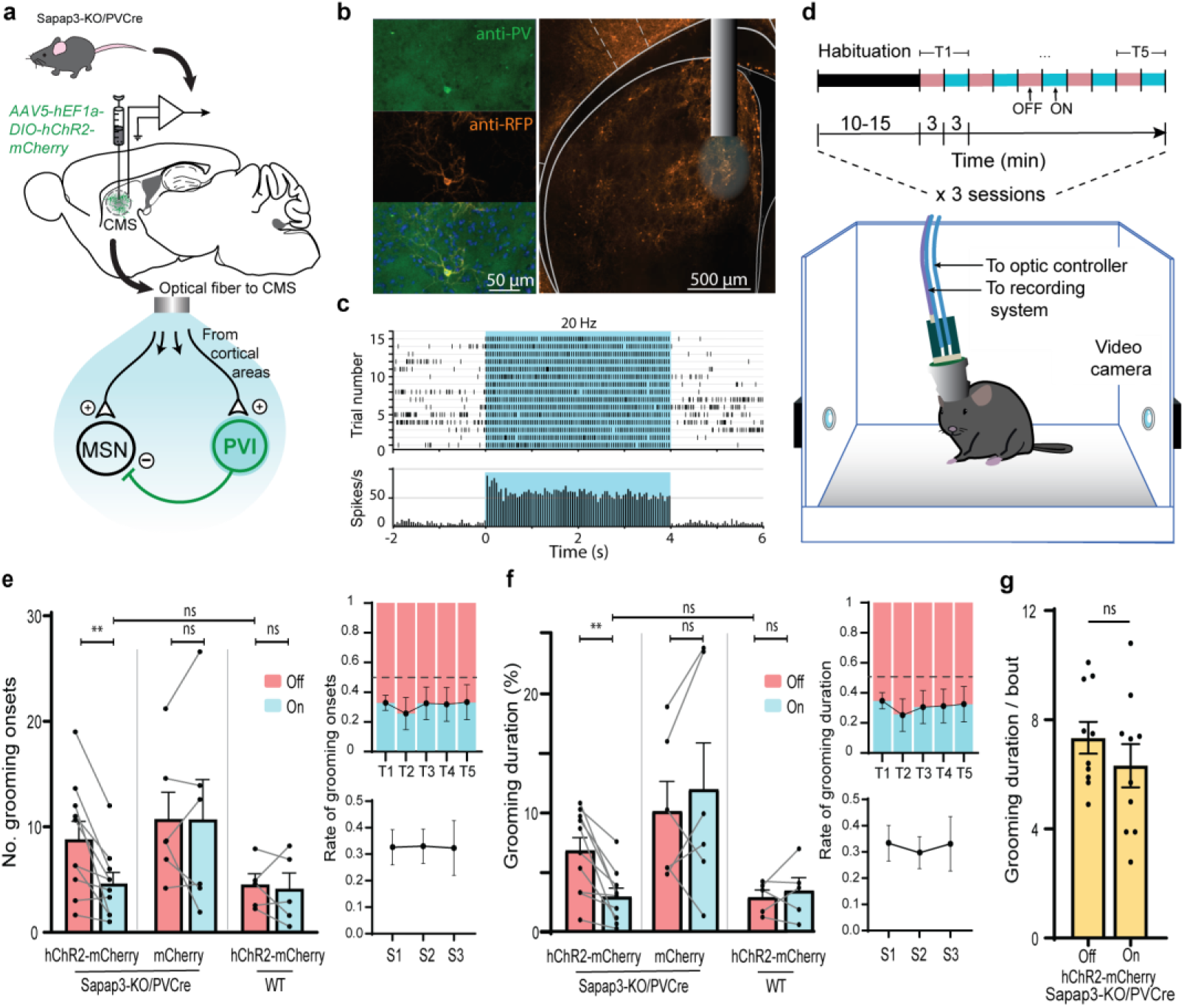
Optogenetic activation of centromedial striatal parvalbumin-positive interneurons reduces compulsive-like behaviour in Sapap3-KO mice. (**a**) Schematic illustration of the experimental design and simplified model of the bilateral centromedial striatal optogenetic neuromodulation of parvalbumin-positive interneuron (PVI) micro circuitry. CMS, centromedial striatum; MSN, medium spiny neuron. (**b**) Representative *post-hoc* histological image illustrating CMS optic fiber placement and specific opsin expression in PVI. RFP, red fluorescent protein. (**c**) Spike raster plot (top) and peri-stimulus time histogram (bottom) confirming optogenetic recruitment of an hChR2-expressing PVI during the light stimulation period (cyan rectangles) over resting state. (**d**) Experimental setup and optogenetic stimulation paradigm with alternating 3-minute blocks of OFF (red bars) and ON (cyan bars) stimulation (20 HZ, 5-ms, 10 mW) trials (T). (**e-f**) Optogenetic activation of CMS PVI reduces the number of grooming onsets and relative grooming duration in Sapap3-KO/PVCre mice expressing hChR2 (n = 10, Wilcoxon matched-pairs signed-rank tests, \*\**P*_*On-OFF, onsets*_ = 0.004, \*\**P*_*On-OFF, duration*_ = 0.002) to wildtype levels (n = 5) (Mann Whitney test, *P*_*onsets*_ = 0.93, *P*_*duration*_ = 0.78). The same CMS PVI injection and stimulation protocol did not alter grooming behaviour in Sapap3-KO/PVCre mice injected with a control virus, only expressing the fluorophore marker mCherry (n = 6, Wilcoxon matched-pairs signed-rank test, *P*_*On-OFF, onsets*_ = 1, *P*_*On-OFF, duration*_ = 0.31), nor in wild-type mice (n = 5, Wilcoxon matched-pairs signed-rank test, *P*_*On-OFF, onsets*_ = 0.81, *P*_*On-OFF, duration*_ = 0.81). Optogenetic treatment was effective across trials and sessions (right panels; GLMM with Poisson distribution and trial, session and mouse as random effects: \*\**P*_ON/OFF, onsets_ = 0.009; \*\*\*\**P*_ON/OFF, duration_ < 0.0001). (**g**) Optogenetic stimulation did not affect the average duration of individual grooming events in Sapap3-KO/PVCre mice (n= 10, Wilcoxon matched-pairs signed-rank tests, *P* = 0.21). Data are presented as mean values ±SEM; ** *P* ≤ 0.01; ns = non-significant.

### II. A transient lOFC delta band signature predicts compulsive-like grooming onsets

To causally prove that striatal PVI recruitment is sufficient for GO regulation, we sought a predictive biomarker to drive their activity before GO. To this end, we focused on one of the most prominent candidate regions implicated in compulsive-like behaviours, the orbitofrontal cortex (OFC)^3,24–26^. Its rodent homolog has been shown to play a crucial neurophysiological role also in the Sapap3-KO mouse model of compulsive-like behaviours^6,9,27,28^. We investigated grooming-related local field potential (LFP) activity in freely moving Sapap3-KO/PVCre mice (n=10) by performing *in-vivo* extracellular tetrode recordings in the lateral orbitofrontal cortex (lOFC). We manually annotated video frames of GO, which we defined as the first noticeable movement of front limbs engaging in a grooming event (Extended Data Fig. 4a). We selected grooming events with no preceding motor behaviours to exclude movement confounders. We contrasted the power spectral density of average pre-grooming activity (-1s to GO) with average resting activity (Fig. 2a). Despite the comparable stillness displayed by the animal during resting and pre-grooming states, their corresponding power spectrum distributions differed, particularly within the delta frequency band (1.5-3 Hz) (Wilcoxon matched-pairs signed-rank test, *P* = 0.01). This low-frequency band power in pre-grooming activity consistently increased towards GO (Fig. 2b). This ramping effect was characterised by an average rising point approximately 1s before GO and an average maximal power of around 2 Hz (Fig. 2c-e). The narrow temporal power increase in delta range frequency preceding GO was observed in individual Sapap3-KO/PVCre mice (Extended Data Fig. 4b). As the average rising point occurred at approximately 1s prior to GO, this power increment embedded the predictive dimension required for driving PVI activity before GO by using closed-loop neuromodulatory intervention. To that end, we developed a supervised learning method for the automatic processing of neural signals acquired via multiple tetrodes (Fig. 3a). The methodological challenge was designing a computationally light LFP signal processing for immediate feedback stimulation. Thus, our technical innovation consisted of decomposing unprocessed signals into a small representative matrix of coefficients reflecting the energy distribution of a given period. This was achieved by implementing triangular filter sets centred on the frequency band of interest, which allowed for capturing small fluctuations around a frequency band of interest while integrating broader information from neighbouring frequency bands. This filter reduction step was essential to train and operate a light feed-forward artificial neural network with an output layer of two outcomes, one for pre-grooming events and one for any other behaviour (OB), such as scratching, rearing, walking, standing, and resting. Given the previously identified GO LFP signature, we designed a filter distribution for each mouse in the low-frequency range between 1-13 Hz (f-max mean= 10±3.27) with seven filters. Taking advantage of such LFP filter decomposition, we drastically reduced the number of input variables for training to 0.01% of the original number of inputs, i.e. by continuously processing one second of 32 acquisition channels sampled at 20 kHz, we reduced the input data dimension to only 35 features. LED stimulation was triggered if LFP activity recorded on at least 50% of the electrodes was classified as pre-grooming. With such an approach, our closed-loop system achieved above-chance performance to predict grooming activity on a trial-by-trail basis for 10.800 decisions per mouse (12 trials of three minutes each, with the algorithm taking a decision every 200 ms) (Fig. 3b).

**Figure 2.**
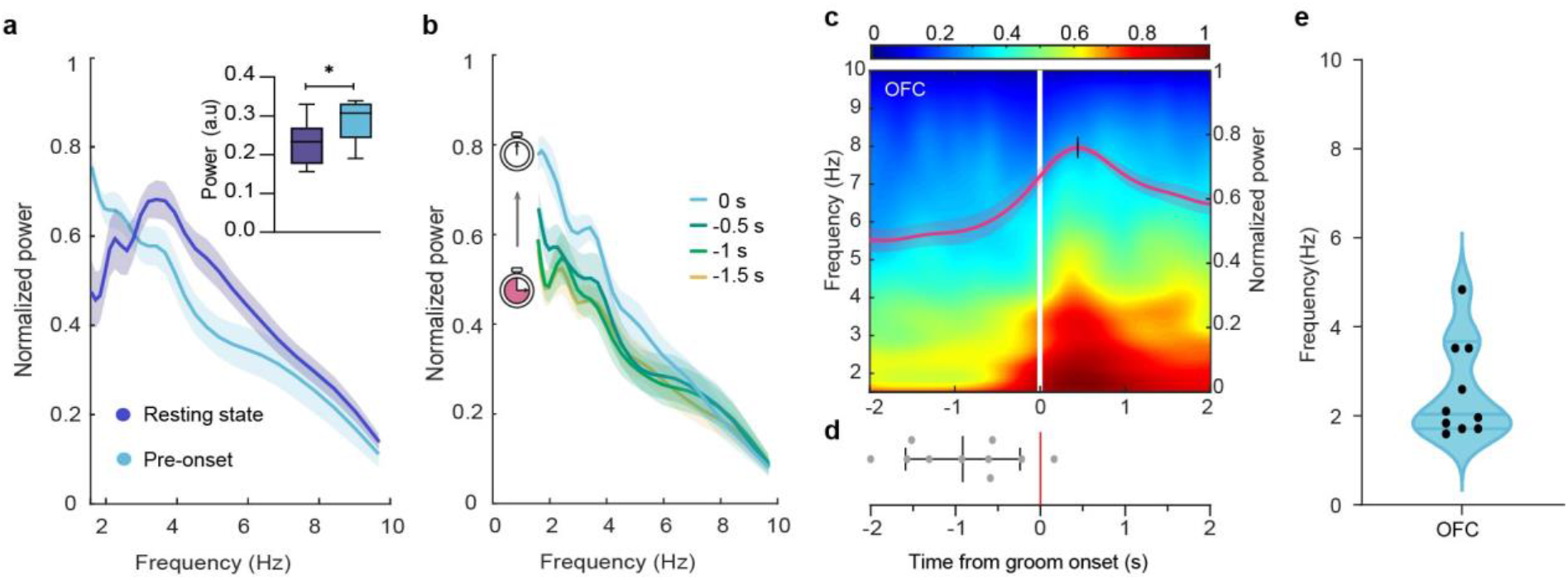
Low-frequency LFP signature in the orbitofrontal cortex predicts grooming onset in Sapap3-KO mice. (**a**) Average power spectral density of OFC LFPs recorded during one-second periods before grooming onset and resting-state in Sapap3-KO mice. The insert illustrates that average OFC LFP power in the 1.5-3 Hz range during pre-grooming exceeds that of the resting state during the same period (n = 15 events for each condition per animal; Wilcoxon matched-pairs signed-rank test, \**P* = 0.01; whiskers represent min-max values). (**b**) Time course of average OFC power spectral density in incrementing 500ms time windows prior to grooming onset of Sapap3-KO mice. (**c**) The average spectrogram for continuous wavelet transform of OFC activity in Sapap3-KO mice displayed a progressive low-frequency power increase before grooming onset (vertical white line). The superimposed red trace represents the normalised power of the average 1.5-4 Hz frequency band (right-hand y-axis). (**d**) Power rising points of the 1.5-4 Hz time-frequency curve are observed on average one second before grooming onset (*M*=-0.914s, *SD*=0.67). (**e**) Frequency values at maximal power are centred on average around 2Hz within 4s peri-grooming onset. Envelopes in a-c represent ±SEM; data in d represent mean ±SD, and data in e represent median and interquartile range. \**P* ≤ 0.05. Each graph depicts the results of n=10 Sapap3-KO mice.

**Figure 3.**
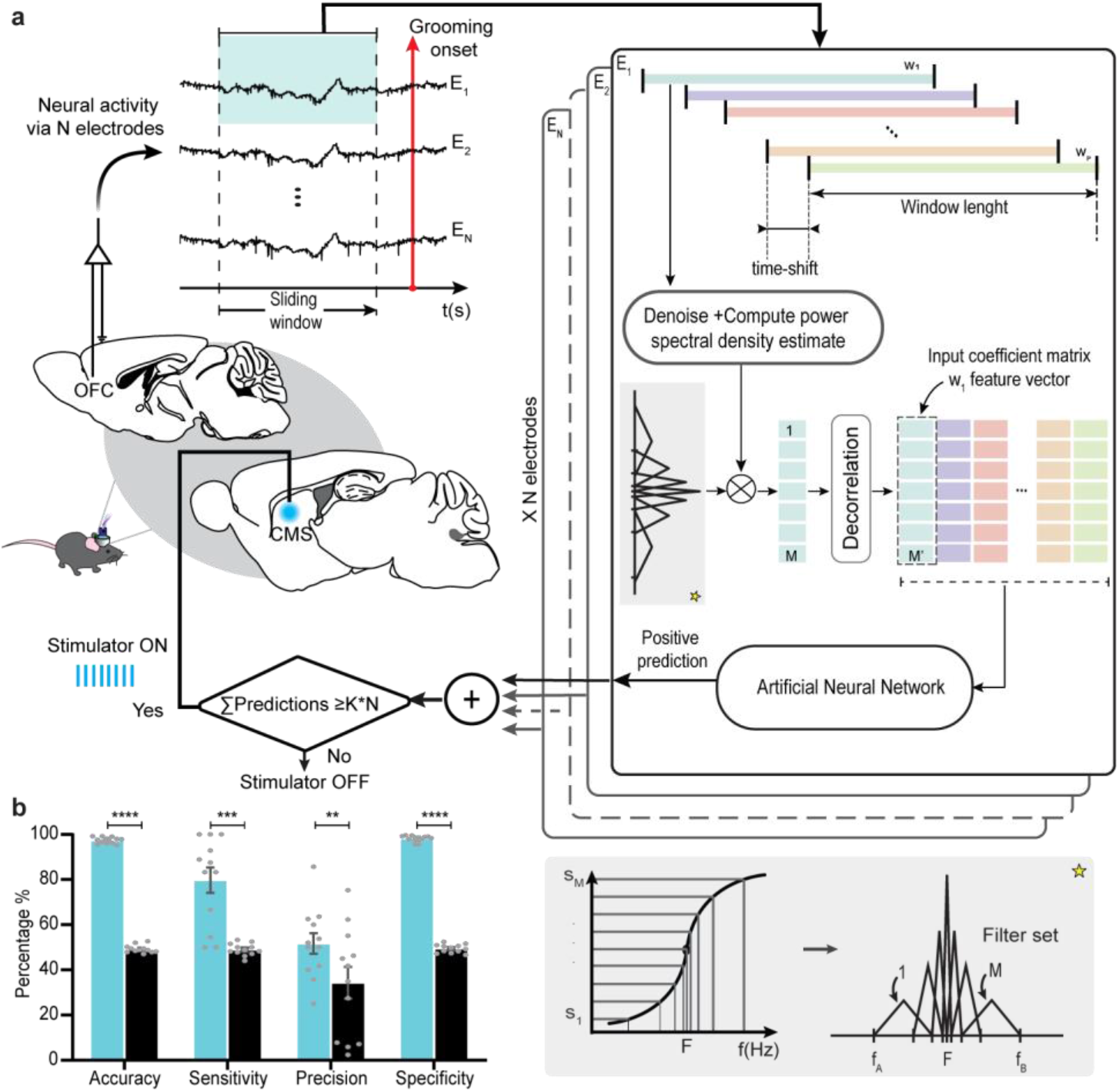
Orbitofrontal cortex low-frequency LFP signature predicts grooming onset in SAPAP3-KO mice. **(a)** Schematic representation of the closed-loop algorithm developed to detect pre-grooming activity online from multiple OFC LFP recording channels. According to the designed pre-processing feature reduction stage, a set of filters is pre-defined around a principal frequency of interest, allowing the detection of small changes around any frequency of interest *F* while integrating broader information from neighbour frequency bands (yellow star, see details in bottom panel). The power spectral density estimate is calculated for each analysis window, and the set of filters is applied to obtain a vector of *M* coefficients. After *P* iterations, it is possible to generate a small matrix of *MxP* decorrelated coefficients that reflect the desired energy distribution in a period. This matrix constituted the input to feed a feed-forward artificial neural network with two output classes, i.e. ‘pre-grooming’ and ‘other behaviour’. This approach enabled the representation of the energy distribution envelope with an input information drastically reduced by 99.99%, allowing real-time signal processing. (**b**) The algorithm’s average classification metrics during online trials (n = 10.800 decisions per mouse, i.e. 12 trials of three minutes each, with the algorithm taking a decision every 200 ms) were contrasted to that of a uniform pseudo-random distribution algorithm with a binary response with the same response rate. All classification metrics were superior for the closed-loop algorithm for accuracy (Paired t-test, \*\*\*\**P*<0.0001), sensitivity (Paired t-test, \*\*\**P*=0.0003), precision (Paired t-test, \*\**P*=0.0018), and specificity (Paired t-test, \*\*\*\**P*<0.0001). Data are presented as mean values ±SEM.

### III. Closed-loop optogenetic stimulation of striatal PVI prevents grooming onset

Taking advantage of the real-time detection of LFP signal in the lOFC predicting grooming onset, we designed a closed-loop approach in Sapap3-KO/PVCre mice expressing hChR2 (H134R)-mCherry. In this approach, the real-time prediction of GO would trigger an optogenetic controller to deliver on-demand light stimulation bilaterally in the CMS to specifically recruit striatal PVI when GO is predicted (Fig. 4a). We designed a 3D-printed homemade headstage for this experimental paradigm that allowed us to implant tetrodes in the lOFC and optic fibers bilaterally in the CMS (Extended Data Fig.2 c-e). Similar to the continuous stimulation procedure, the experimental paradigm included a habituation stage of 10-15 minutes, followed by alternating 3-minute trials of ‘Off’, ‘Closed-loop’, and ‘Yoked’ stimulation (12 trials total) (Fig. 4b). During ‘Off’ trials, no stimulation was delivered. During ‘Closed-loop’ trials, grooming events were automatically predicted online, which triggered the delivery of blue light pulses for 4 seconds (5-ms pulses at 20Hz and 10mW). During ‘Yoked’ trials, the number of light stimulations equalled the preceding ‘Closed-loop’ trial; however, they were randomly assigned across the 3 minutes. This protocol was repeated on three different days. We found that closed-loop optogenetic stimulation significantly reduced GO in Sapap3-KO/PVCre mice (n=5) compared to ‘Off’ and ‘Yoked’ trials (Fig. 4c). As in the continuous stimulation protocol, this effect was consistent across trials and sessions (Extended Data Fig. 5) and observed across all animals (Fig. 4c, bottom). The number of bouts and the grooming duration were reduced by 59.37% and 70.54%, respectively (Fig. 4d, e and Extended Data Fig. 6a). In line with results obtained in continuous optogenetic stimulation experiments, the character and duration of individual grooming sequences remained unaltered (Extended Data Fig. 6b). Finally, our ‘closed-loop’ approach was as efficient as the ‘continuous’ stimulation protocol in terms of reducing the number of grooming bouts (Fig. 4d) and grooming duration (Fig. 4e), but with a tremendous reduction of stimulation time by 87.2% (Fig. 4f).

**Figure 4.**
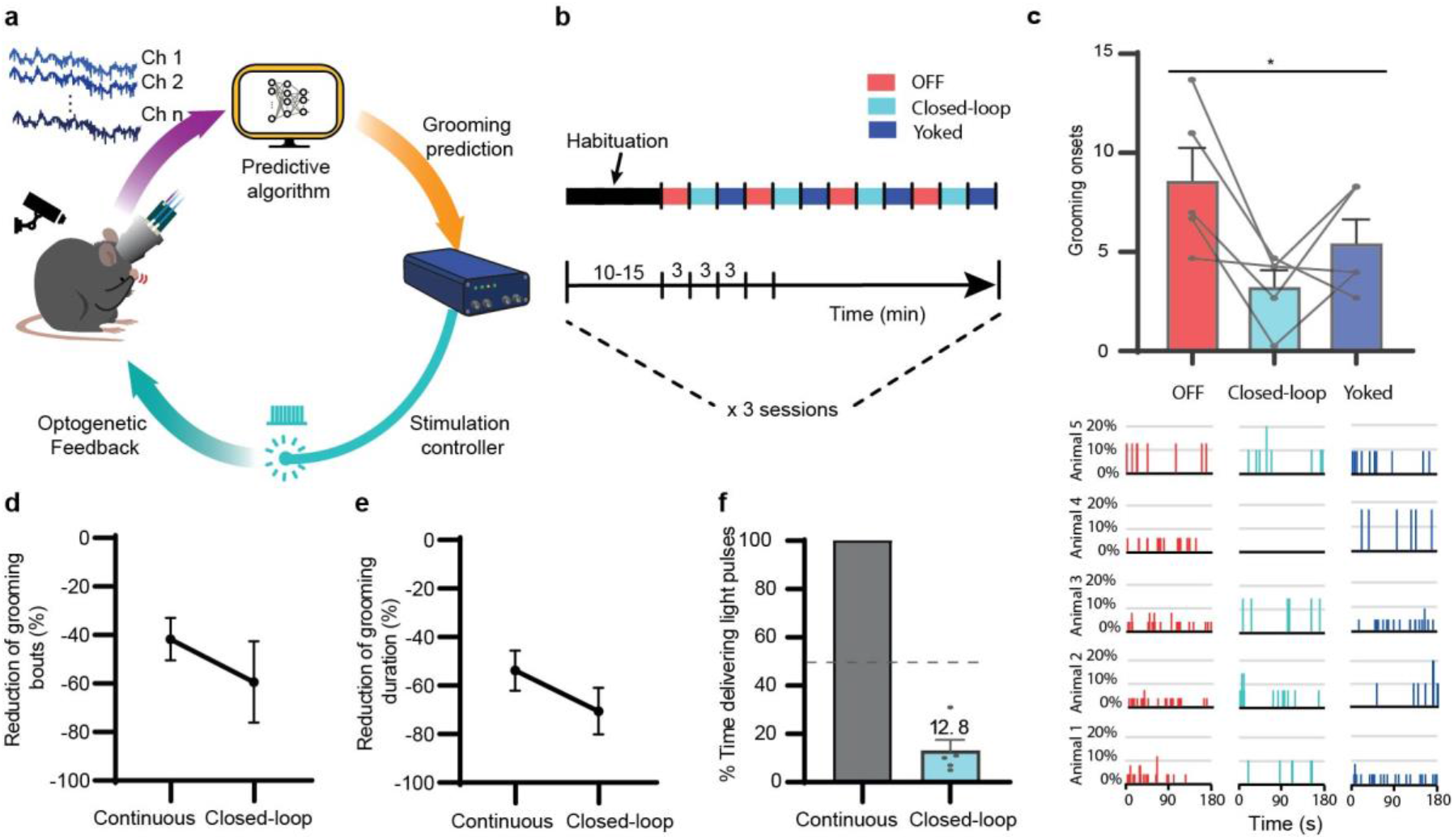
Closed-loop optogenetic intervention alleviates compulsive grooming in Sapap3-KO/PVCre mice. **(a)** Schematic representation of the closed-loop experimental paradigm for on-demand optogenetic intervention. OFC LFPs are acquired in Sapap3-KO/PVCre mice via chronic implants for neural activity recording and light stimulation. OFC LFPs are digitalised and processed using the novel online feature decomposition and reduction methods described above (purple arrow). If the supervised classification algorithm detected the low-frequency biomarker predictive of grooming onset through categorising the processed OCF LFPs, a control signal (orange arrow) is sent to activate a stimulation controller to deliver blue light pulses during 4s. (**b**) Illustration of the closed-loop optogenetic stimulation protocol alternating “OFF” (red), closed-loop” ON” (cyan) and “yoked” (blue) stimulation trials. (**c**) The closed-loop optogenetic stimulation reduced the average number of grooming onsets of Sapap3-KO/PVCre relative to OFF and yoked trials (top panel) (Page’s L test for ordered alternatives, H=Closed-loop<Yoked<OFF, \**P*=0.02; GLMM, \**P*=0.02). The corresponding histograms for each animal display the distribution of grooming onsets across the experimental period (bottom panels). Histograms for each animal display the distribution of grooming onsets across the experimental period for each experimental condition (bottom panel). (**d**) Percentage of reduction in grooming bouts and (**e**) in grooming duration for continuous and closed-loop conditions. (**f**) Comparison of relative stimulation time between continuous and closed-loop stimulation protocols. Data represent means ±SEM.; * *P* ≤ 0.05. Each graph depicts results of n=5 Sapap3-KO/PVCre mice.

## DISCUSSION

By combining optogenetic neuromodulation with a closed-loop stimulation protocol, we demonstrated in our study that the recruitment of striatal PVI prior to grooming onset was sufficient to regulate compulsive-like behaviours of the Sapap3-KO mice. We first showed that continuous optogenetic stimulation of the PVI network in the centromedial striatum immediately reduced the number of compulsive-like grooming events in Sapap3-KO mice to the level of matched wild-type controls. Next, when seeking a neurophysiological predictor of compulsive-like events to define the critical period for the recruitment of the PVI network, we detected an increase in the spectral power of the delta band activity in the lateral OFC, which shortly and consistently preceded grooming onset. To our knowledge, this is the first report of transient LFP events in the OFC that predict compulsive-like grooming episodes. Using this predictor, we designed a closed-loop approach based on an innovative supervised machine learning protocol to recruit the striatal PVI network prior to compulsive-like grooming onset. Via this closed-loop system, we showed that brief optogenetic PVI excitation upon grooming event prediction was sufficient to abort their occurrence and, as a consequence, to reduce compulsive-like behaviours in Sapap3-KO mice. Taken together, we believe that these results will be valuable for understanding the neurobiological mechanisms underlying repetitive behaviours; furthermore, they also open up new alleyways for designing therapeutic interventions for the treatment of pathological repetitive behaviours.

Indeed, our results showed that striatal PVI are essential in preventing the onset of compulsive-like behaviours in Sapap3^-/-^ mice, presumably by inhibiting hyper-active MSN projection neurons^6,19^. These observations align with previous studies that showed the implication of striatal PVI during the suppression of prepotent inappropriate actions, regulation of choice execution, or generation of complex motor sequences^29–31^. The crucial role of striatal PVI in regulating compulsive-like grooming in the Sapap3-KO mice is also supported by recent studies describing abnormal synaptic properties of striatal PVI in this animal model^32^ as well as by observations in other mouse models suffering from pathological repetitive behaviours where abnormally low PVI density in striatal areas was observed^22^. Our study proposes that the temporal recruitment of striatal PVI shortly before the onset of compulsive-like behaviours is crucial to regulating their initiation.

However, the mechanisms by which the striatal PVI network could be recruited still need to be characterised. Interestingly, two recent studies have shown a coincidence in the emergence of delta oscillations in the prefrontal cortex and the recruitment of pyramidal cell assemblies^33,34^. In the context of our study, it would be interesting to investigate whether such recruitment of pyramidal cell assemblies occurs through the here observed OFC delta oscillations, which preceded compulsive-like grooming. Consequently, such cortical drive could allow for the temporally adjusted recruitment of the striatal PVI network, which in turn regulates the expression of compulsive-like grooming behaviours. This hypothesis is supported by the observation of a decreased spectral power in the low-frequency band in the OFC of Sapap3-KO compared to wild-type mice, including in the delta range^27^. Thus, a lower LFP delta band in the OFC of Sapap3-KO mice may underlie impaired recruitment of downstream striatal PVI networks and, thus, a decreased regulation of compulsive-like behaviours. Our findings also echo a recent study in OCD patients implanted with recording-capable DBS devices; the authors showed that LFP signals recorded in the striatum were co-occurring with OCD symptoms and detected a signature of interest in the delta-band, as we observed in our study^35^.

Our findings also call for testing potential therapeutic strategies where striatal PVI could be specifically targeted in pathologies with compulsive behaviours. For instance, electrical deep brain stimulation (DBS) protocols have been used in severe OCD patients with good but limited efficiency in reducing compulsive symptoms while targeting different brain structures along with basal ganglia circuits^36–38^. As continuous electrical DBS does not differentiate between different neuronal populations, this therapeutic approach offers room for improvement by adapting stimulation protocols to reach cell-type specificity. Such a possibility has been confirmed in a recent study where the authors demonstrated that they could specifically recruit PVI in the globus pallidus by using brief bursts of electrical stimulation^39^. Building upon these findings and our results, it will be interesting to test such optimised DBS procedures in the context of pathological repetitive behaviours in order to recruit striatal PVI specifically.

Another interesting aspect of our study was developing an innovative supervised machine learning approach for the automated delivery of on-demand optogenetic stimulation. The technical innovation of our approach for continuous processing of electrophysiological brain signals was to develop a decomposition stage for the drastic reduction of online-handled variables. Thus, the following signal processing stages require less computing power and allow for simpler architectures that could be implemented in general-purpose computers, as we did in our experiments. We believe that this methodology would be of great interest for exploiting neurophysiological signals in a real-time manner, such as adaptive DBS and, more generally, in brain-machine interface processes^40,41^. Our closed-loop approach was as efficient in decreasing compulsive-like grooming as our continuous approach, with a reduction of the stimulation time by 87%, a crucial parameter in DBS therapy for battery saving. Even more interesting than this technical advantage is investigating potential long-term therapeutic effects such as recently observed using patterned neuromodulation protocols^39^. Finally, we believed that the implemented predictive approach might serve as a platform to further explore other brain targets for therapeutic purposes in a closed-loop manner.

## FIGURES

**Extended Data Figure 1.**
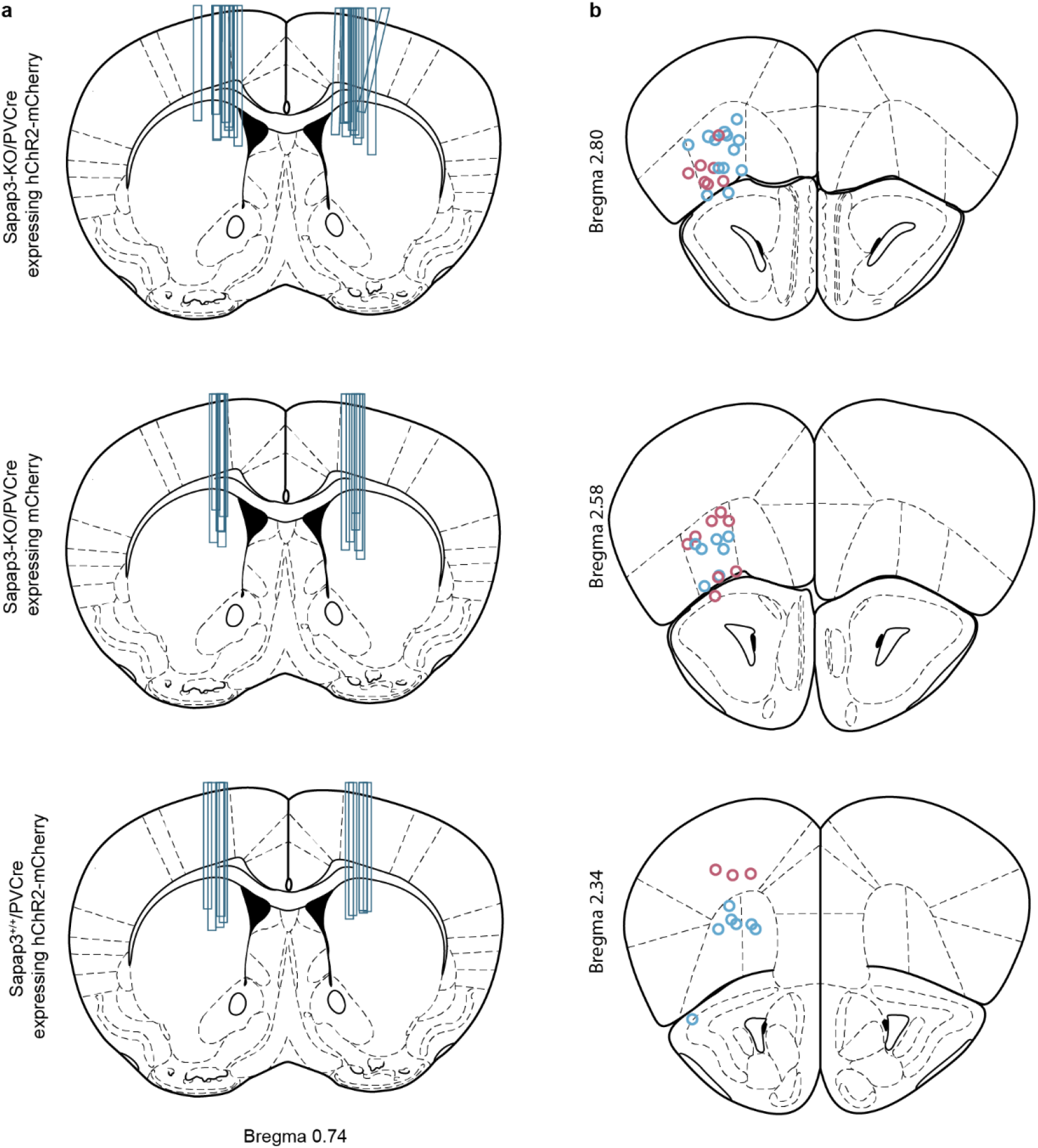
*Post-hoc* histological confirmation of optic fiber and tetrode tip placements in CMS and lOFC. (**a**) Reconstruction of individual fiber placements in the CMS of Sapap3-KO/PV-Cre mice expressing hChR2-mCherry (upper panel), of Sapap3-KO/PV-Cre mice expressing mCherry (centre panel), and of Sapap3^+/+^/PVCre mice expressing hChR2-mCherry (bottom panel). The location of each fiber tip was confirmed by comparing fiber tracks in histological coronal sections with measured fiber lengths of the detached implant after perfusion of the implanted animals. (**b**) Reconstruction of individual tetrode tips as electrophysiological recording sites in the lOFC, mapped to coronal sections of a mouse brain atlas (Paxinos, G. & Franklin, K. B. J., 2008). The location of each tetrode tip was determined based on electrolytic mark lesions in prefrontal coronal sections or measuring of tetrode lengths of the detached implant post perfusion. All illustrated recording sites were used for the LFP signature (n = 10 Sapap3-KO/PV-Cre mice); recording sites, which were additionally used for closed-loop stimulation experiments are depicted in blue (n = 5 Sapap3-KO/PV-Cre mice). CMS = centromedial striatum; lOFC = lateral orbitofrontal cortex.

**Extended Data Figure 2.**
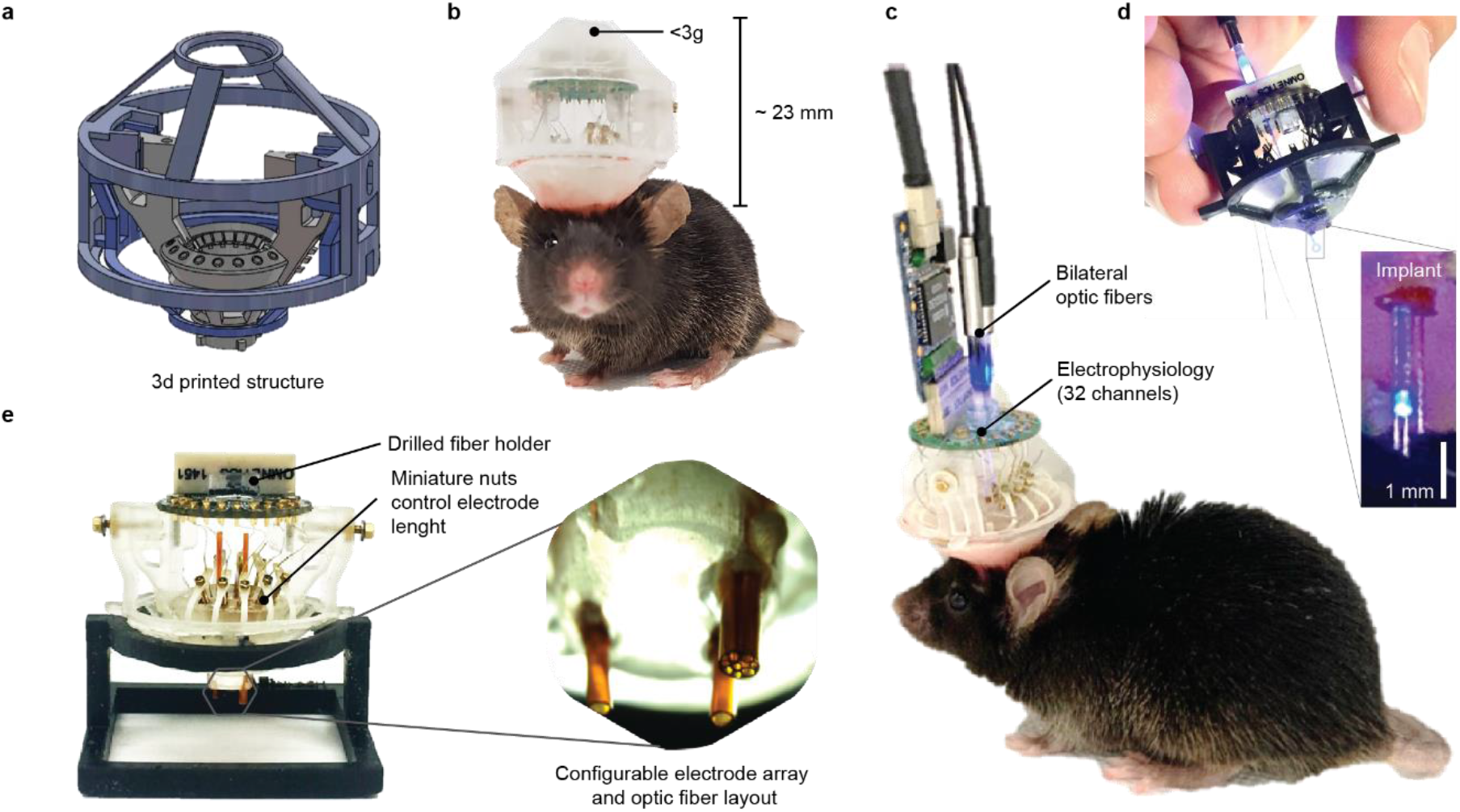
Cranial implant design for chronic electrophysiological recordings and optogenetic neuromodulation in freely moving mice. (**a**) Isometric view of the 3D model structure depicting the three individual printable pieces that assemble the implant: the lower protection cone, the removable protection cap and the element for adjusting tetrode positions (‘drive’), which is a custom-adapted version of the “flexDrive” model^42^. (**b**) Picture of a mouse implanted with our custom-built chronic device, including the protection cap. (**c**) Picture of a chronically implanted mouse tethered for electrophysiology activity recording and optogenetic stimulation. (**d**) Picture of the implant during testing of optical power through optic fibers. Close-up view of an extracted implant after experimental procedures, showing one striatal fiber and surrounding tetrodes for optotagging of recorded PVIs. (**e**) Picture of the custom-adapted configurable 3D printed driving mechanism targeting multiple brain regions and allowing for the adjustment of individual tetrode depths for optimising chronic electrophysiological recordings over the duration of several weeks. Close-up bottom view of the implant showing the bottom end of the tubing inside of which either optical fibers or up to eight tetrodes can be placed.

**Extended Data Figure 3.**
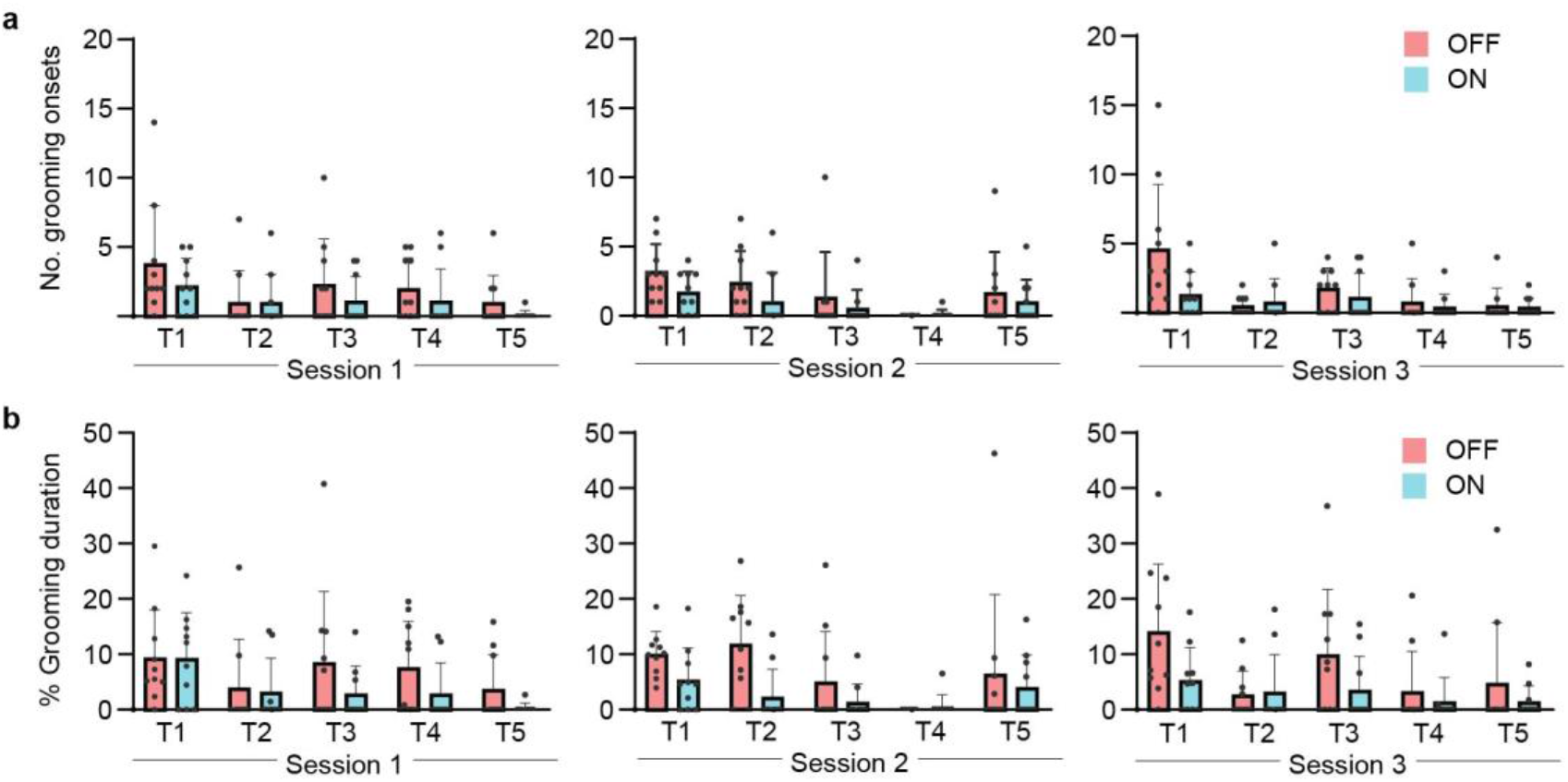
Detailed behavioural results of optogenetic stimulation of centromedial striatal PV interneurons in Sapap3-KO/PVCre mice. (**a**) Number of grooming onsets and (**b**) the percentage of grooming duration across individual consecutive trial blocks (T1-T5) and three separate sessions (1-3). Optogenetic treatment was effective across trials and sessions (right panels; GLMM with Poisson distribution and trial, session and mouse as random effects: \*\**P*_ON/OFF, onsets_ = 0.009; \*\*\*\**P*_ON/OFF, duration_ < 0.0001). Each graph depicts n=10 Sapap3-KO/PVCre mice results, and the data are presented as mean values +SEM.

**Extended Data Figure 4.**
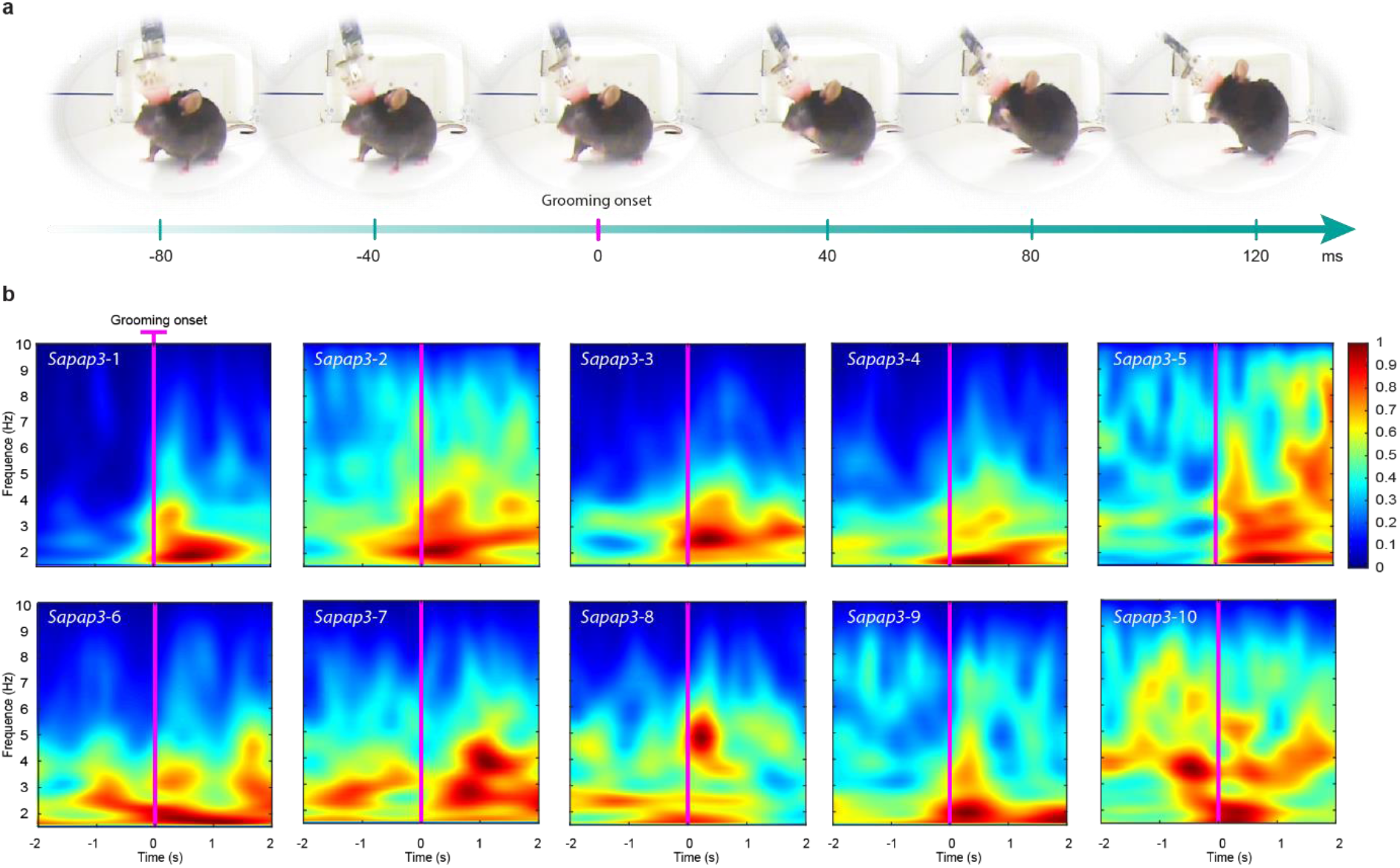
Low-frequency LFP signature in the OFC predicts grooming onset. (**a**) Example of consecutive image frames displaying grooming onset and the assignment of the exact time point labelled “grooming onset” to the first frame showing the upwards locomotion of the anterior limbs towards the orofacial area to engage in a grooming bout. (**b**) Individual time-frequency wavelet spectrogram around grooming onset (-2 to +2 seconds peri-grooming onset) reveals an increase in low-frequency power in each recorded animal (n=10 Sapap3-KO mice). Vertical magenta lines in each spectrogram indicate grooming onset as assigned in panel a. Each spectrogram represents the average normalised LFP activity of grooming events (*M* = 25 events, *SD* ±5.25) for a single mouse extracted over different recording sessions; warmer colours represent higher energy.

**Extended Data Figure 5.**
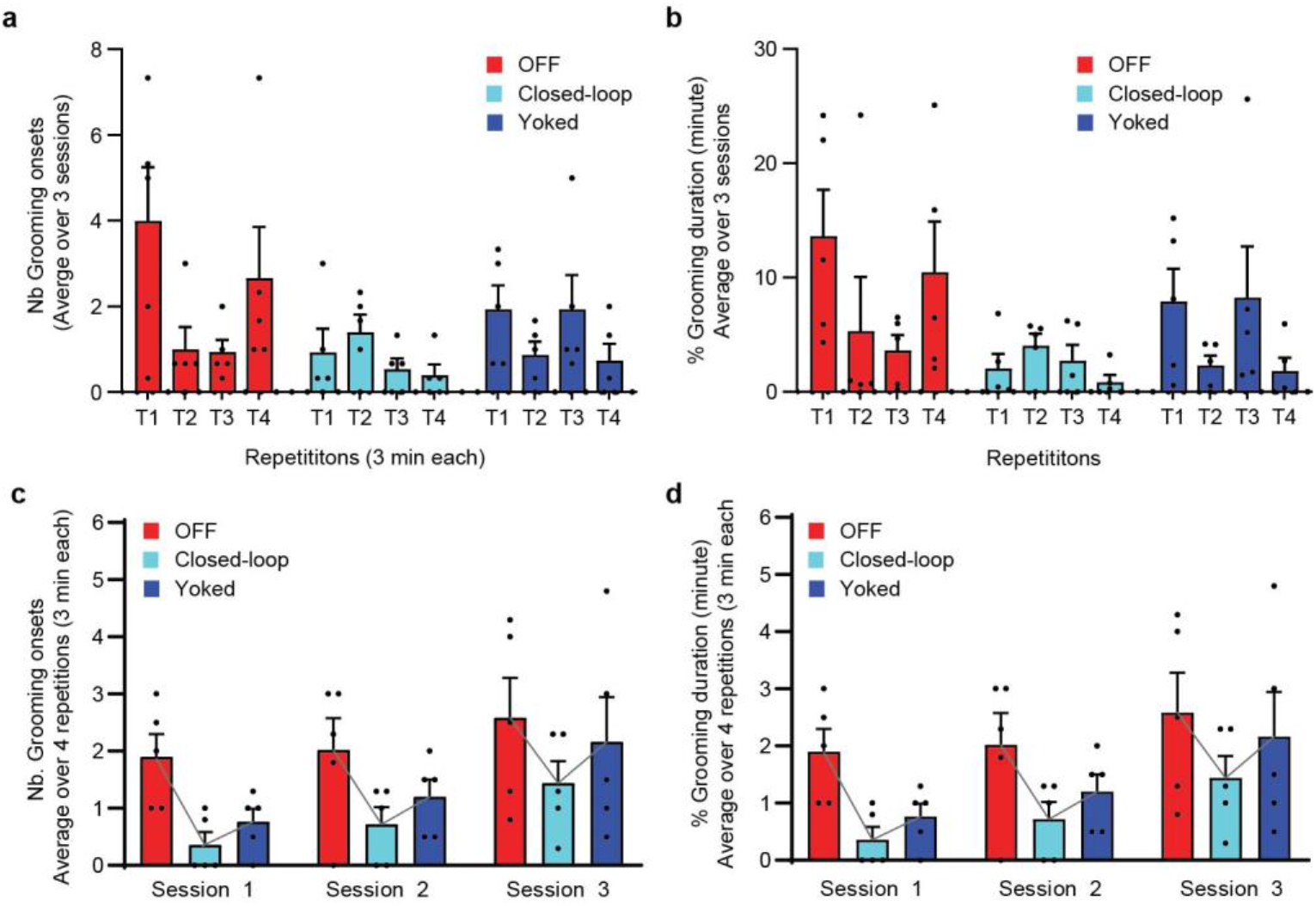
Intra- and inter-session comparison of OFF, closed-loop, and yoked optogenetic stimulation. (**a**) Detailed overview of the number of grooming onsets and the (**b**) percentage of grooming duration, segregated into individual repeated trials (T1-T4) of OFF (red), Closed-loop ON (cyan) and Yoked (blue) optogenetic stimulation. Each dot represents the average of each individual mouse across three experimental sessions. (**c**) Detailed overview of the number of grooming onsets and the (**d**) percentage of grooming duration, segregated into sessions (1-3) and OFF (red), Closed-loop ON (cyan), and yoked (blue) optogenetic stimulation condition. Individual dots represent the average of each individual mouse across four different trials per session. All graphs depict the behaviour of n = 5 Sapap3-KO/PVCre mice. Data are presented as mean values +SEM. The grey line in panels (**c**) and (**d**) highlights the trend of average grooming number or onsets across the three stimulation protocols. A significant effect of the stimulation condition was found while integrating the variables trials (a,b) and sessions (c,d) along with the subject as random effect factors; this was the case both for the number of grooming onsets (GLMM, \**P*< 0.01) and for the percentage of grooming duration (GLMM, \*\**P*< 0.01).

**Extended Data Figure 6.**
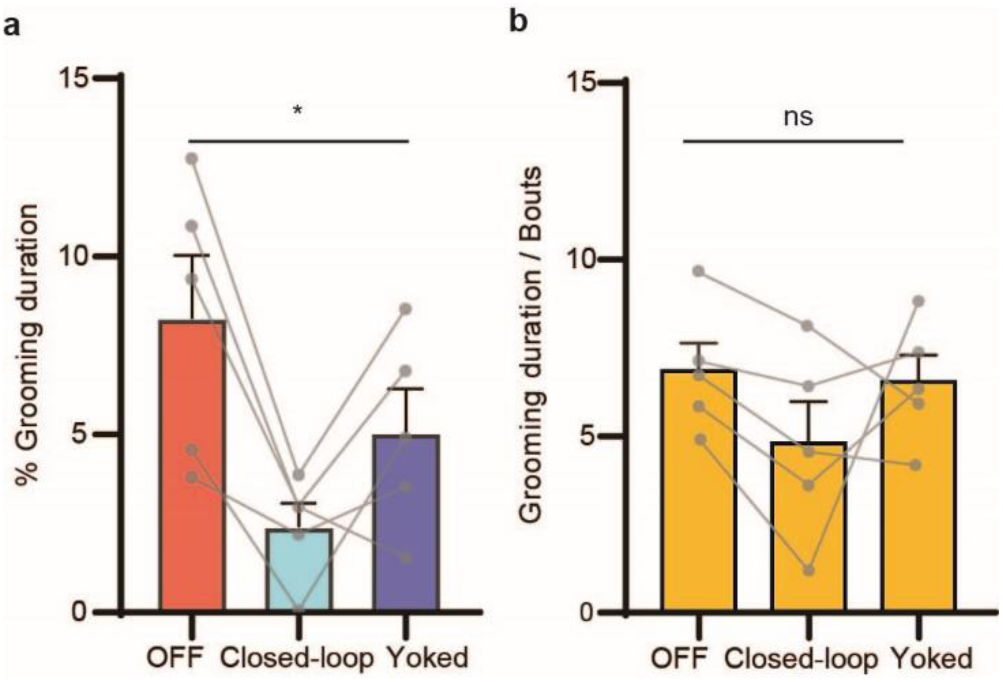
Closed-loop optogenetic stimulation effect on grooming duration. (**a**) Closed-loop optogenetic activation of CMS PVI reduced the average percentage of grooming duration in Sapap3-KO/PVCre compared to “OFF” and “Yoked” trials (Page’s L test for ordered alternatives, H=Closed-loop<Yoked<OFF, L=68, \**P*=0.01; GLMM, \*\**P*=0.002). Grey dots and interconnecting lines represent the average of individual animals across trials and sessions. (**b**) The stimulation protocol did not affect the average duration of individual grooming events in either the “OFF”, “Closed-loop”, or the ‘Yoked’ condition (repeated measures one-way ANOVA, *P* = 0.9). Data are depicted as means (+SEM) in the n=5 Sapap3-KO/PVCre mice tested in the protocol.

## METHODS

### Mice

All experimental procedures have been approved by the French Ministry of Higher Education, Research and Innovation (APAFIS #1418-2015120217347265 and #31141-2021042017105235). Mice were maintained in a 12-hour light/dark cycle with *ad libitum* food and water. The Sapap3-knockout (Sapap3-KO) mouse line was provided by Drs. G. Feng and A.M. Graybiel (Massachusetts Institute of Technology, Cambridge, U.S.A.) (B6.129-*Dlgap3*^*tm1Gfng*^/J ; Jackson Laboratory stock #008733). *Sapap3*^+/−^ were crossed with parvalbumin (PV) (*Pvalb*)-Cre line (B6;129P2-*Pvalb*^*tm1(cre)Arbr*^/J; Jackson Laboratory stock #008069; provided by Dr A. Bacci, Paris Brain Institute, France) to obtain Cre-positive *Sapap3*^+/+^ and Sapap3-KO mice. Biopsies for genotyping were taken during weaning and at the end of all experimental procedures to verify the absence or presence of the Sapap3 and Cre-recombinase proteins, respectively, according to genotyping recommendations of the Jackson Laboratory (standard PCR assay protocols #24678 and #27783 for PV-Cre and Sapap3-KO line, respectively). For experimental procedures, we used 17 male Sapap3^-/-^ :: PV-^Cre/wt^ (average initial age = 8.6 ±1.4 months, average weight = 30.5 ±3.7 g) and 5 age-matched Sapap3^+/+^::PV-^Cre/wt^ (average initial age = 9.1 ±1 months, average weight = 39.3 ±4 g).

### Viral Vectors

Viral vectors expressing either an excitatory opsin and a fluorophore reporter (AAV5-hEF1a-dlox-hChR2(134R)-mCherry(rev)-dlox-WPREhGHpA) or a fluorophore reporter only (AAV5-hSyn1-dlox-mCherry(rev)-dlox-WPRE-hGHpA) were purchased from the Viral Vector facilities of the Neuroscience Center Zurich (University of Zurich and ETH Zurich) and Addgene Viral Service facilities. Viral vectors were generated from Addgene plasmids #20297 and #50459, and titers were determined via fluorometric quantification (Neuroscience Center Zurich) or by real-time quantitative PCR combined with SYBR green technology (Addgene). Respective titers were >9.1×10^**12**^ vector genomes/ml (Neuroscience Center Zurich, opsin construct), >1.3×10^**13**^ vector genomes/ml (Neuroscience Center Zurich, control construct) and 1.4 × 10^13^ GC/mL (Addgene, opsin construct). Until stereotaxic injection surgeries, the viral vectors were aliquoted and stored at -80°C until stereotaxic injection surgeries.

### Headframe implant

Homemade implants were designed and constructed around a transparent plastic core, a modified version of the Open Ephy’s flexdrive^42^ (Extended Data Figure 2a). The implant was designed with SolidWorks 3D CAD software (Dassault Systems SolidWorks Corporation, MA, USA) and printed via stereolithography using the 3D printer Form2 (Formlabs, Somerville, MA, USA) with clear resin. The plastic drive body core held eight independently mobile tetrodes and two fixed flat optic fiber stubs (Plexon Inc., Texas, USA) of 200/230 μm (core/core + cladding) inserted in a zirconia ferrule (Extended Data Figure 2c-e). Tetrodes were made from 17.78 μm (0.0007 inches) Formvar-coated nichrome wire (A-M Systems, WA, USA), twisted (Tetrode Assembly Station, Neuralynx, MT, USA), and heated form tetrodes. Tetrodes were fixed to laser-cut plastic springs (polyethylene terephthalate, 25 mm from Weber Metaux, Paris, France) and individually driven via miniature screws (0.7M Micro-modele, Strasbourg, France). Electrodes were attached to an electrode interface board (Open Ephys Production Site, Lisbon, Portugal) with gold pins (Neuralynx, MT, USA). Individual electrodes were gold plated to an impedance of 200-350 kΩ. Ground connections were made using 50.8 μm (0.002 inches) Formvar-coated tungsten wires (A-M Systems, WA, USA). The fully assembled implant (drive body, cap, and cone) weighed less than 3 g.

### Surgical procedures

Each mouse received an injection of analgesics (buprenorphine 0.1mg/kg; s.c.) 30 minutes prior to deep anaesthesia (induction at 2.5% % isoflurane; maintenance during surgery at 0.9-1.2% isoflurane). The animals were gently mounted into a digitally equipped stereotaxic frame (David Kopf Instruments) via ear bars adapted for mouse stereotaxic surgeries. The animal’s temperature was maintained at 37°C through a heating pad coupled to an anal sonde. Scalp hair was removed using depilation cream (Veet) diluted with sterile 0.9% saline, and the skin was disinfected three times with 70% ethanol and betadine solution (Vétédine, Vétoquinol) before skin incision. Skull and skin were maintained humid during the entire surgery using sterile 0.9% saline. Four small burr holes were drilled around the perimeter of the exposed skull surface to accept steel anchor screws. Two additional small burr holes were drilled behind the estimated implant volume to insert two separate ground wires. For all mice, craniotomies and durotomies (1.0mm diameter) were made bilaterally above the dorsomedial striatum (AP = +1.0 mm, ML = +1.5 mm); for mice with *in-vivo* tetrode recordings in the orbitofrontal cortex (OFC), an additional cranio-/durotomy (0.8mm diameter) was performed above the left lateral orbitofrontal cortex (AP = +2.8 mm, ML = +1.5 mm). The brain surface was maintained humid with sterile 0.9% saline throughout viral injections until headframe implantation. AAV constructs were injected bilaterally at a constant rate of 50nl/min (0.4μl/site) into the dorsomedial striatum (DV -2.4, measured from the brain surface) using a motorised micro pump (Legato 130, KD Scientific) with a precision syringe (Hamilton Gastight Series #1701, 10μl) and respective needles (Hamilton, 33Ga, bevelled end). Before virus injection, the needle was lowered to DV = -2.5 mm and retracted to DV = -2.4 mm immediately, where the needle was allowed to settle for 2 minutes. After injections, the needle was maintained in the same position for 10 minutes, then retracted by 300μm (DV = -2.1) and maintained at that position for another minute. Afterwards, the needle was slowly removed from the brain. After each injection, to exclude needle clogging during intracranial procedures, correct flux was tested by visually checking the continuous formation of a droplet using the same injection speed as during intracranial injections. A drop of surgical lubricant was applied to each cranial opening. The implant was carefully lowered so that the bottom polyamide tubes containing either tetrodes or smaller tubing to hold the optic fibers touched the brain surface. The implant was fixed to the skull and screws with dental acrylic (Jet Denture Repair, Fibred Pink powder + liquid, Lang Dental). Ground wires were formed into tiny loops inserted below the skull to make surface contact with the brain. Ground wires were then fixed to the implant frame and connected to the electrode interface board (EIB). Optic fibers were carefully lowered to the depth of 2mm into the brain and glued into position in the opening provided in the EIB. Tetrodes were advanced individually directly after surgery in the orbitofrontal cortex (DV = 1.0 mm) or the striatum (DV = 1.5 - 2 mm) during the following 3-5 postoperative days. A protective 3D-printed cap was screwed to the implant to keep out bedding debris. Animals were injected with analgesics (buprenorphine, 0.1mg/kg; s.c.) directly after surgery and every 12 hours post-surgery on the following day; they were closely monitored until awake in a heating chamber set to 37°C. Animals were then placed into a clean home cage equipped with materials adapted for implanted animals (e.g. no hanging food grids to avoid bumping the implant). Animals were closely monitored post-surgery twice a day, including veterinary care, taking into account changes in weight, body score, nesting, and overall locomotor activity as indicators of surgery recovery.

### Experimental setup

The experimental setup consisted of a homemade arena (290 × 270 mm) in which the mouse was allowed to move freely. The setup structure included passive commutators for electrophysiology and optic stimulation, synchronisation LEDs, and two cameras (704×576 resolution; 25 frames per second) installed at opposite walls to allow for complementary views. After surgical recovery, the animals were habituated for three days to human handling, the electrophysiology acquisition system and the fiber patch cable tethering. The animals were left in the arena with *ad-libitum* food and water for two nights (6 pm to 8 am) of additional habituation to the setup. At the beginning of each experimental session, the mice were carefully tethered and habituated to the arena for 10-15 minutes. Any noises and vibrations were avoided during experimentation. No water or food was provided during the experimental sessions. After each experiment, the behavioural apparatus was cleaned using a disinfectant cleaning spray containing 55% ethanol, washed with soap and water, and dried.

### Behavioural assessment

Manuel scoring was performed using a freely available video scoring software (Kinovea, 0.8.15, www.kinovea.org). Self-grooming activity was assessed by quantifying duration, number of grooming events, and the percentage of grooming time. Grooming onset was defined as when the mouse started lifting its front paws to groom. The end of grooming was defined as the time point when the mouse stopped grooming for at least one second or when the grooming behaviour was interrupted by another behaviour. We discarded grooming-like phases of less than a second duration. We annotated other behaviours such as, e.g. walking, rearing, sniffing, resting, stretching, heading up, or freezing. The neural activity during these behaviours was used to form a database for our supervised learning algorithm. We scored resting behaviours for the local field potential analysis; these were defined as behavioural activity without major head movements. Hereby the head position was either resting on the floor or lifted as a relaxed, horizontal extension of the spine. The four limbs touched the ground, and the mouse showed a relaxed posture. I.e. no hunched posture or exaggerated respiration as observed for freezing behaviour was stated; ears were not pulled back, and eyes normally opened, not squinted as observed during distress.

### Optogenetics

Bilateral optogenetic neuromodulation in the dorsomedial striatum was conducted at least two weeks after stereotaxic viral injections to allow for sufficient recovery and viral expression. The implanted striatal fiber stubs were connected to optical patch cables (200-μm core and 0.5 m length with 0.66 numerical aperture) via mating ferrules in a zirconia sleeve. Optical patch cables were connected to blue light LED source modules (465 nm) mounted on magnetic LED commutators. The light stimulation patterns were pre-programmed using Radiant Software (Plexon Inc.). Before experimentation, the output power was measured from the optical fiber tips with a light power meter (Thorlabs PM100D with S120C sensor) and calibrated to 10 mW. Unless otherwise stated, all devices for optogenetic modulations were purchased from Plexon Inc. (Texas, USA).

#### Continuous optogenetic stimulation

Each experimental session lasted 30 minutes and included ten interleaved OFF (no light stimulation) and ten ON trials (light stimulation of 10mW during 5ms at pulses of 20Hz) of three minutes each.

#### Closed-loop optogenetic stimulation

Each experimental session lasted 36 minutes. In four repetitive cycles, we alternated OFF trials, trials with closed-loop stimulation (CL) and with yoked stimulation (Y), each lasting 3 minutes. A custom Matlab (R2017b) algorithm triggered the light stimulation on demand for the CL trials. Each activation lasted four seconds (10 mW, 5-ms pulses at 20 Hz). In a Y trial, the same number of stimulations delivered in the previous CL trial were pre-assigned randomly (*rand* function) by the algorithm.

### Data acquisition and analyses

#### Electrophysiology

We performed extracellular recordings in awake, freely moving mice using tetrodes attached to a 32-channel connector in a mechanically adjustable drive. All signals were amplified, multiplexed and digitalised at a sampling frequency of 20 kHz using the Intan hardware acquisition system (RHD2000 USB Interface Board with the RHD 32-Channel Headstage, Intan Technologies, CA, USA) and either Intan Software (RHD USB Interface Board software) or Open Ephys G.U.I.^43^. Recordings were analysed offline. We used a custom Matlab code to record with Intan Libraries for the closed-loop experiments. A blinking LED, and a digital input activated simultaneously were used to synchronise video and extracellular recordings.

#### Local field potential analysis

Neural signals were synchronised with video recordings via TTL signals for local field potential recordings. The spectral content of the LFP signals was analysed using custom Matlab routines, the Chronux^44^ Matlab package (http://chronux.org) and the Matlab Signal Processing toolbox. For Figure 2 and Extended Data Figure 4, we decreased the broadband data sample rate by 40 and then low-pass filtered the data using a 10th-order Butterworth filter with a cut off frequency of 10 Hz. We performed continuous wavelet transformation around grooming events (4s before and after the onset). We used the Morse wavelet with asymmetry parameter (γ) equal to 3 and a time-bandwidth product equal to 60.

#### Spike analysis

Spike sorting was performed offline using the valley-seeking prevalent method in Offline sorter (Version 3.3.5, Plexon Inc.), followed by visual screening. Each set of spike clusters was compared for cross-correlogram features and spike waveform. The spikes of each putative unit were assessed qualitatively regarding ISI-waveform, spike amplitude and waveform consistency. To quantify the effect of light, we used NeuroExplorer (Version 4, Nex Technologies, Colorado, USA.) rate histogram to display the firing rate versus time with a bin size of 0.05s

### Feature selection and supervised classification algorithm

Extracellular signals in the OFC from freely moving mice were acquired through 32 chronic drive implant recording channels (20 kHz sampling frequency), amplified and digitalised. To obtain the predictive value of low-frequency LFP components in the OFC related to the animal’s behavioural state (“pre-grooming” and “other behaviour”), we designed a pre-processing procedure for feature reduction that captures small changes in the energy distribution in a particular frequency band over time. A set of triangular filters pre-defined by a symmetrical distribution around the principal frequency of interest F (Fig. 2e) within the range from *f*_a_ to *f*_b_ is chosen to represent the shape and distribution of energy of the LFP biomarker in the frequency domain (in our case *f*_a_ = 1, *f*_b_ = 10±3.27, N= 5). The filter distribution is configured to have gradually decaying information concerning the portion of the spectrum to which they are applied. Since the features extracted depend on the distribution of the filter set, the distribution function and its sampling are suited to detect a variety of neural signatures for which a change in energy at a specific frequency band is observed. For each analysis window (sub-segments of one second with 200 ms time-shift), we calculate the power spectral density estimate, and the set of filters is applied to obtain a vector of *M* coefficients whose values are furthermore decorrelated using the discrete cosine transformation. After *P* iterations, we create a matrix of *MxP* decorrelated coefficients (*M=7, P=5*) that reflect the desired energy distribution in a period. This matrix constituted the input for an artificial neural network with a minimal architecture: one input layer with 35 neurons, two hidden layers, and an output layer with two neurons, one for pre-compulsive behaviour and the other for a collection of different types of behaviour. In this setup, one decision is made every 200 ms. We created a database of pre-grooming events and other behaviours for each mouse to train the algorithm. These databases were created from five recording sessions (40 min each), each containing a minimum of 100 events. The database was randomly split into a training data set (70%) and a test dataset (30%). The training datasets were extracted and shuffled from five one-hour recording sessions for each experimental animal on different days. Each electrode was processed individually. We set a threshold policy to decide if the output values of the neural network corresponded to a pre-grooming event or not; only if more than 50% of these detections pointed to a pre-grooming event it was counted as such. When put into practice in the closed-loop experiments, one positive outcome of pre-grooming classification would trigger the optogenetic stimulator for four seconds, a period in which the algorithm would not make any further decisions. For each outcome of the “other behaviour” class, the stimulator would remain off, and the tests would continue every 200 ms.

To estimate the algorithm’s performance in a real-time scenario, we tested the algorithm within a closed-loop protocol of 3 days (Fig. 4a-b), a total of 12 3-minute trials in a Sapap3-/-:: PV-Cre/wt injected with a viral vector expressing only a fluorophore reporter (AAV5-hSyn1-dlox-mCherry(rev)-dlox-WPRE-hGHpA), i.e. in a total of n = 10800 decisions that the algorithm performed. We defined the “early detection period” (*Ep*) as the time window between -2s to grooming onset to half-duration of the grooming event. True positive classifications were the predictions or light stimulations that fell within the *Ep*. False positives were all the other stimulations that did not fall inside the *Ep*. True negative results are the other behaviour predictions outside *Ep* and grooming events, and false negatives are all missed grooming events where no grooming prediction falls into the *Ep*. To contrast our results with that of a pseudo-random classifier, we used the behavioural data collected in the previous closed-loop experiments (12 trials) and implemented a uniform pseudo-random number generator (1 for grooming and 0 for other behaviour) with the same result rate (200 ms). We calculated the accuracy, precision, sensitivity, and specificity for both algorithms and each 3-minute trial.

### Histology

After experimental procedures, electrolytic mark lesions were made to confirm the localisation of tetrode recording sites. Animals were anaesthetised as described above, and 5μA constant current was delivered to each electrode for 20 s using an isolated current stimulator (DS3, Digitimer Ltd, Hertfordshire, England). After 72 hours, mice were anaesthetised via an intraperitoneal injection of pentobarbital (200mg/kg) and transcardially perfused with 30 ml of 4°C cold 0.9% sodium chloride solution followed by 60ml of 4°C cold 4% PFA in 0.1M PB. Brains were post-fixed in the same paraformaldehyde solution overnight at 4°C, briefly rinsed three times in 1xPBS, and progressively dehydrated for cryosectioning by incubations in 15% and subsequently in 30% sucrose solution in 1xPBS for 24 and 48 hours, respectively. Next, brains were embedded in the OCT compound and sectioned into six series of 40μm coronal sections (Microm HM N°560, Thermo Scientific) into 1xPBS containing 0.1% sodium azide. Prefrontal sections were stained with a freshly filtered 1% cresyl violet solution and analysed for the location of marked lesions under a brightfield microscope. Immunofluorescence stainings were performed on the striatal sections of one series in 4°C cold solutions on an orbital shaker. Concretely, sections were washed three times for ten minutes in 1xPBS, followed by three washes of each ten minutes in 1x PBS containing 0.1% Tween 20 and 0.2% Triton-X. Sections were next blocked for two hours in 5% normal goat serum in 1xPBS containing 0.1% Tween 20 and 0.2% Triton-X. Subsequently, sections were incubated in the same blocking buffer containing anti-red fluorescent protein antibody (polyclonal rabbit anti-RFP, Rockland, #600-401-379, Lot #35634; dilution: 1:1000) on an orbital shaker at 4°C overnight. Sections were then washed three times for ten minutes in 1xPBS with 0.1% Tween 20 and 0.2% Triton-X and incubated for two hours in a blocking buffer containing a secondary antibody (polyclonal anti-rabbit Cy3-conjugated AffiniPure, produced in goat, Lot 106489; dilution 1:400). Afterwards, sections were washed twice for ten minutes in 1xPBS with 0.1% Tween 20 and 0.2% Triton-X, followed by one washing for ten minutes in 1xPBS. Next, sections were incubated for 2 minutes and 30 seconds in DAPI solution (10μg/ml), washed three times for ten minutes in 1xPBS and mounted in 0.1M phosphate buffer onto Superfrost Plus slides and coverslipped using a fluorescence-protecting medium (FluoromountTM, Sigma Aldrich). A selection of sections was incubated in the same buffer with an additional anti-parvalbumin antibody (polyclonal guinea pig anti-PV GP 72, Swant, dilution 1:5000) in order to verify viral infection of specifically parvalbumin-positive cells. The secondary antibody solution additionally contained polyclonal anti-guinea pig Alexa488, produced in goats (Invitrogen, Lot #145863, dilution: 1:400). All striatal sections were imaged using a slide scanner (Axio Scan.Z1, ZEISS), and the sections were analysed and mapped using ZEN software (ZEISS) and a mouse brain atlas^45^.

### Statistical analyses

All statistical analyses using R were conducted using R version 4.1.1 (R Development Core Team, 2021) and those using Prism were conducted using version 8.0.1 (GraphPad Software Inc). All levels of statistical significance were set at p-values < 0.05.

#### Continuous stimulation analyses

We used Prism to perform the following statistical analyses for analyses of continuous ON-OFF experiments: Wilcoxon matched-pairs signed-rank test (Paired, non-parametric test, two-tailed, confidence level 95%); Mann Whitney test (Unpaired, non-parametric test, comparing ranks, two-tailed, confidence level 95%); and one-way ANOVA. To assess the potential effects of the factors “trial” or “session”, we performed an Aligned rank transform ANOVA including an additional mouse as a random effect, using the package ARTool in R version 4.1.1 (R Development Core Team, 2021). Lastly, a generalised linear mixed model using the *Poisson* family was fitted to explain the “number of grooming” by the different “treatments”. Hereby, the model included “MouseID”, “Trial” and “Session” as random effects. “Trial” random effect was nested to “Session” random effect. The significance of the main effects of continuous ON/OFF stimulation (“treatment”) was assessed based on Type II Wald chi-square tests. Please note that the *“glmmTMB”* function (from *glmmTMB* package) was used to model the GLMM because it is intended to handle zero inflation, unlike the *glmer* function. Finally, we fitted a linear mixed model to explain also the grooming duration by continuous ON/OFF stimulation (“treatment”). Again, the model included MouseID, Trial and Day as random effects. The trial random effect was nested to day random effect. The square-root transformation was used on the grooming duration response variable to improve the model assumptions of linearity, normality, and constant variance of residuals. The significance of the main effects of ON/OFF treatment was assessed based on Type II Wald chi-square tests.

#### Analyses of the classification algorithm

To assess the performance of our supervised classification algorithm compared to a pseudo-random classification algorithm, we applied paired t-tests for each performance parameter, i.e. accuracy, sensitivity, precision, and specificity, after prior verification of the normal distribution of the data.

#### Closed-loop stimulation analyses

To assess the statistical evidence for the increase in the ordinal ranks between three “treatments” (ClosedLoop, Yoked and Off) in terms of the number of grooming, the repeated measure trend test, *Page’s L test* for ordered alternatives ^46^ was used with the *PageTest* R function in *DescTools* package where the ordered alternative hypothesis H1, m_ClosedLoop < m_Yoked < m_Off, was tested against the null hypothesis H0, m_ClosedLoop = m_Yoked = m_Off. Hereby, values were averages by animal and session. Additionally, A generalised linear mixed model using the *Poisson* family was fitted to explain the “number of grooming” by the different stimulation “Treatments” (Closed Loop, Yoked and OFF). The model included MouseID, Trial and Day as random effects. The random trial effect was nested to day random effect. The significance of the main effects of Treatments was assessed based on Type II Wald chi-square tests. Post hoc pairwise comparisons were carried out on a significant Treatment effect with the *“emmeans”* package to determine where the differences occurred across the treatments. P-values resulting from the post hoc tests were adjusted to control for the “false discovery rate” (FDR) due to multiple comparisons. We also conducted a *Page’s L test* for ordered alternatives as described above to assess the significance of stimulation conditions on grooming duration.

Additionally, we fitted a GLMM to explain grooming duration by the different stimulations (Closed Loop, Yoked, Off). The model included MouseID, Trial and Day as random effects. The random trial effect was nested to day random effect. The square-root transformation was used on grooming duration as a response variable to improve the model assumptions of linearity, normality, and constant variance of residuals. The significance of the main effects of treatment was assessed based on Type II Wald chi-square tests. Post hoc pairwise comparisons were carried out on a significant Treatment effect with the *“emmeans”* package to determine where the differences occurred across the Treatments. P-values resulting from the post hoc tests were adjusted to control the “false discovery rate” (FDR) due to multiple comparisons. Lastly, to compare average grooming bout durations in the Off, Closed-loop, and Yoked stimulation conditions, we performed a one-way ANOVA using Prism (GraphPad version 8.0.1).

## Data availability

The data supporting this study’s findings are available from the corresponding author upon reasonable request.

## Code availability

The custom code for the LFP analysis, the feature selection, and the supervised learning algorithm supporting this study’s findings are available from the corresponding author upon reasonable request.

## Author Contributions

Conceptualisation and supervision: EB; software development: SLMG; hardware development: SLMG, EB, CS; colony management: SLMG, CS; experimental work: SLMG, CS; data analysis: SLMG, CS, EB; writing of the manuscript: SLMG, CS, EB; graphical illustration: SLMG, CS, EB.

## Acknowledgements

All animal work was conducted at the PHENO-ICMice facility. This work also benefitted from the equipment and services from the iGenSeq core facility at ICM for the genotyping of the animals, from the equipment and advice by the ICM Histomics core facility for histological validation, from the equipment and advice by the ICM Workshop core facility, as well as from statistical advice and analyses by Sana Rebbah and François-Xavier Lejeune from the ICM Data Analysis core facility. We furthermore thank Pr. Jean-Luc Zarader for advice on the pre-processing and feature extraction of the closed-loop algorithm. This work was realised with the following funding: Agence Nationale de la Recherche (ANR-16-INSERM-SINREP, ANR-19-ICM-DOPALOOPS) (EB); Fondation de France (EB); Carnot Foundation (EB); the Science and Technology Council of Mexico (CONACYT) (SLMG); ANR-*Investissement d’Avenir* program (Labex Bio-Psy) (EB, SLMG); and the L’Oréal-UNESCO Fellowship 2016 (CS). The core facilities were supported by “Investissements d’avenir” (ANR-10-IAIHU-06 and ANR-11-INBS-0011-NeurATRIS) and “Fondation pour la Recherche Médicale”.

